# Identifying Gait Phases from Joint Kinematics during Walking with Switched Linear Dynamical Systems

**DOI:** 10.1101/378380

**Authors:** Luke Drnach, Irfan Essa, Lena H. Ting

**Affiliations:** Georgia Institute of Technology, Atlanta, GA, 30332 USA; Emory University, Atlanta, GA, 30322 USA

## Abstract

Human-robot interaction (HRI) for gait rehabilitation could benefit from data-driven, subject-specific gait models that account for gait phases and gait dynamics. Here we address the current limitation in gait models driven by averaged kinematic data, which do not model interlimb gait dynamics and have not been shown to precisely identify gait events. We used Switched Linear Dynamical Systems (SLDS) to model joint angle kinematic data from healthy individuals walking on a treadmill during normal gait and during gait perturbed by electrical muscle stimulation. We compared model-inferred gait phases to gait phases measured independently via a force plate. We found that SLDS models accounted for over 88% of the variation in each joint angle and labeled the joint kinematics with the correct gait phase with 84% precision on average. The transitions between hidden states matched measured gait events, with a median absolute difference of 25ms. To our knowledge, this is the first time that SLDS inferred gait phases have been validated by an external measure of gait, instead of against pre-defined gait phase durations. SLDS provide individual-specific representations of gait that incorporate both gait phases and gait dynamics. SLDS may be useful for developing control policies for HRI aimed at improving gait by allowing for changes in control to be precisely timed to different gait phases.

## I. INTRODUCTION

Developing human-robot interactions (HRI) for gait assistance and rehabilitation could benefit from models of gait phases and gait dynamics that are both data-driven and computable in real-time. Gait phases provide important information about discrete changes in body dynamics that occur with each **gait event**, i.e. ground contact and lift-off of each foot. These gait events define four **gait phases**, as the body dynamics alternate between single- and double-limb support: single limb support is defined as the left and right swing phases, when the trailing limb lifts off the ground at toe-off to when it contacts the ground at heel-strike; double support phases are defined as the periods when both limbs are on the ground between swing phases. The gait phases also define important periods of whole-body stability to external perturbations, which should be considered in physical HRI. In double-limb support, the body has a wider base of support and is more stable with respect to external perturbations. Multijoint coordination patterns and lower limb dynamics are also different within each gait phase, and are likely to differ across individuals or with motor impairment. Individual-specific models of gait dynamics within each gait phase may advance the development of physical HRI strategies that identify and correct deficits in limb coordination during gait. HRI strategies designed for specific gait phases could leverage the differences in whole body stability in single versus double limb support. Here we address the current limitation in data-driven models of gait, which do not model interlimb gait dynamics, have not been shown to precisely identify gait events, and are not subject-specific.

Joint kinematic data has previously been labeled with gait phases using Gaussian Mixture Hidden Markov Models (GHMMs), which infer a series of discrete states from measured data. Previous work using GHMMs to model gait have established that the hidden states have the same duration as the gait phases, both in healthy populations [1] and in populations with pathological gaits [2,3]. However, the correspondence between the GHMM states and gait phase transitions times has not been validated against external measures of gait phase, especially on a step-by-step basis.

GHMM approaches also lack models of gait dynamics necessary for control synthesis and for predicting future states. GHMMs assume the observed kinematics are independent of each other across time and are normally distributed within each phase, creating a static, statistical model of gait kinematics [1,2]. GHMMs can only describe gait in terms of the mean and covariance within each phase; they cannot recreate trajectories of limb motion or simulate the effects of external perturbations on the joint angles. Some authors have attempted to circumvent this problem by applying cubic splines to GHMM-generated samples to reconstruct gait [3]. Although such models may be sufficient for discriminating different gaits or gait phases, it is unclear how they could be used in real-time control, because it would be difficult to model the effect of control inputs on gait dynamics.

Gait dynamics have been modeled using switched linear dynamical systems (SLDS), which identify a set of linear dynamical systems (LDS) with each hidden state of an HMM (Figure 1). SLDS have been used to distinguish gaits such as walking, jogging, and limping [4–6]. The advantage of an SLDS over a GHMM is that the future joint angle trajectories can be predicted if the times for switching between linear models [7] or the conditions on joint kinematics for switching [8] are known. Previous SLDS models of gait have not explicitly represented interlimb dynamics of single and double-support gait phases, instead modeling each limb separately, and included two [7] or four [8] hidden states per leg. Switching conditions were estimated by assuming the gait phase duration of each state. As such, the correspondence between model states and actual gait phase was not tested.

**Fig. 1.**
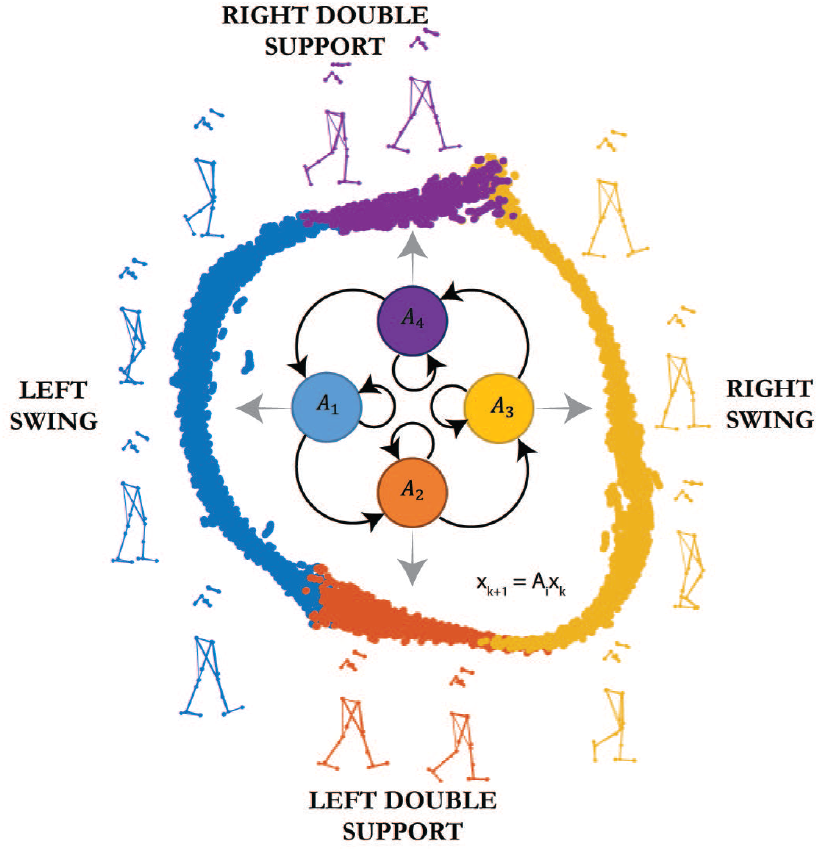
SLDS model of gait. Individual postures are grouped together by kinematic similarity using t-SNE [9], and are color-coded according to the SLDS model hidden state. Stick figures indicate the postures represented by the neighboring points. In the center is a schematic diagram of the four states of the SLDS, and their conditional dependence on each other. The SLDS model learns to segment the gait kinematics according to kinematically similar points, and the assignment corresponds to gait phases.

Here, our objective is to demonstrate that an SLDS model of gait, trained on an individual’s specific gait kinematics, can accurately label kinematic data with gait phases in normal and perturbed gait on a step-by-step basis, and can represent the limb motion dynamics within each gait phase. We focused on developing individual-specific models of gait because, in practice, gait impairments affect each person’s gait differently; as such, the HRI for rehabilitation will need to account for and correct individual differences. We modeled kinematic data from healthy individuals walking on a treadmill in normal conditions. We hypothesized that the four gait phases - left leg swing phase (left swing), double support with left leg forward (left double support), right leg swing phase (right swing), and double support with right leg forward (right double support) - could be modeled as four hidden states using a single LDS for each state. We compared the sequence of hidden states generated by the SLDS to an independent and standard measure of gait phase, obtained via a force plate. We tested the robustness of the model to differences in gait dynamics by assessing the accuracy of the predicted gait phase, using the SLDS from normal gait, on the unmodeled, perturbed gait. Our results indicate that, in both normal and perturbed gait, SLDS hidden states accurately label joint kinematic data with the correct gait phase, and that transitions between the hidden states correspond precisely to gait events.

## II. Methods

### A. Gait Data

We used data collected from five healthy participants walking at constant speed on a treadmill (Table 1). All participants consented to the protocols that were approved by the Emory Institutional Review Board. Hip flexion and adduction, knee flexion, and ankle flexion and adduction angles of both legs were measured at 100Hz using motion capture (Vicon Motion Systems). Force plates in the treadmill recorded ground reaction forces at both feet at 1kHz.

**TABLE I.** Participant Characteristics

| Participant | Sex | Height (cm) | Mass (kg) | Walking Speed (m/s) |
| --- | --- | --- | --- | --- |
| 1 | F | 160 | 63 | 1.0 |
| 2 | F | 174 | 57 | 0.9 |
| 3 | F | 172 | 68 | 0.9 |
| 4 | F | 164 | 59 | 1.1 |
| 5 | F | 173 | 64 | 1.1 |

We used data from baseline walking on a treadmill and perturbed walking with muscle stimulation. The complete experimental protocol consisted of three conditions: baseline walking, perturbed walking, and walking post-perturbation. For each participant, the walking speed was held constant across all conditions, and ranged from 0.9-1.1m/s. In the perturbed walking condition, functional electrical stimulation (FES) was applied to the right dorsiflexors during right swing and to the right plantarflexors during late stance, which resulted in increased right ankle flexion during right swing phase. Participants walked continuously through both perturbation and post-perturbation conditions, with bouts of each condition lasting for 45 seconds resulting in about 45 gait cycles per walking bout. The baseline walking condition occurred at the beginning of the experiment, before any stimulation was applied, and at the end of the experiment, after the last post-perturbation period [10,11].

### B. Switched Linear Dynamical Systems

A SLDS is of a set of LDSs with hidden states that govern when to switch between the individual linear models. The equation for an SLDS in discrete-time can be written as:

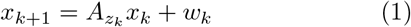

where *x_k_* the observed state at time *k, z_k_* is the hidden state at time *k, A_zk_* is the linear dynamics matrix associated with the hidden state at time k, and w_k_ is the residual term, assumed to be zero-mean and Gaussian, with covariance matrix Σ*_z_k__* determined by the current hidden state. At each instant in time, the hidden state *z_k_* is a one of a finite set of states, *z_k_* ∈ {1, …, *N*}. The probability of starting in each hidden state is given by the initial state distribution, *π*, and transitioning between hidden states is governed by a transition probability:

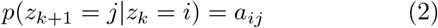

Estimating the model parameters *θ* = (*A, π, Σ, a*) occurs by maximizing the log-likelihood of observing the data, given the model parameters:

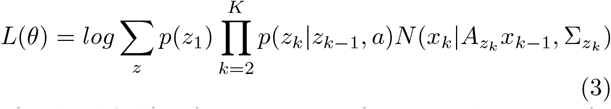

The log-likelihood is maximized iteratively using the Expectation-Maximization algorithm, also called the Baum-Welch algorithm in the context of HMMs. In the expectation step, we calculate the joint probability of transitioning from hidden state *i* to hidden state *j*:

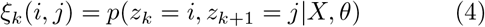

and the posterior probability of being in hidden state *i*:

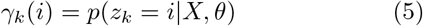

where both probabilities are estimated using the forward-backward procedure. In the maximization step, we update the model parameters to maximize the log-likelihood [12]:

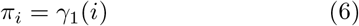

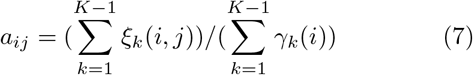

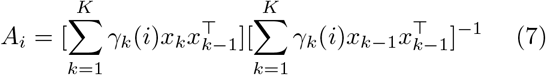

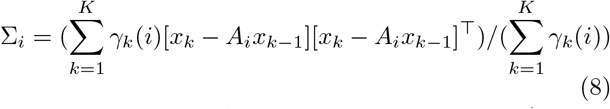

Starting from an initial set of parameters, we iterate between the expectation and maximization steps until the difference in log-likelihood between consecutive iterations was smaller than a predefined threshold value.

### C. Model Training

To capture individual-specific gait dynamics, we trained a four-state SLDS for each individual on unperturbed, constant speed walking. We used the current and previous joint angle as our state vector, 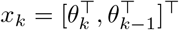, where *θ_k_* is the vector of measured joint angles at time *k*. The transitions between hidden states were constrained to form a cycle, where only self-transitions and transitions to the next state were allowed.

Because the EM algorithm locally maximizes the log-likelihood, the training results for SLDS are highly sensitive to the initial estimates of the linear dynamics. To get an accurate model of gait phase dynamics, we devised an initialization procedure to estimate the linear models from kinematic features. We initialized the gait phase by determining the gait events that mark the transition between the phases. We initialized heel strikes as the time of the minima in knee flexion angle and toe-offs as the time of the minima in ankle flexion angle. Using our kinematically determined gait events, we initialized the linear dynamics coefficients and the prediction error covariances for each LDS in each gait phase using multivariable regression.

To reduce variability in the model parameters due to the small amount of training data, we averaged the model parameters for a single individual over multiple training instances. We divided each participant’s baseline walking data into five equal-length segments, each containing about 9 gait cycles, and repeated initialization and training five times, leaving out one fifth of the data each time. This procedure gave us five estimates of the model parameters, which we averaged together to form the final parameter values. Using the average model parameters, we applied the Viterbi algorithm [13] to determine each person’s model-predicted gait phases in both baseline and perturbed walking conditions (Figure 2).

**Fig. 2.**
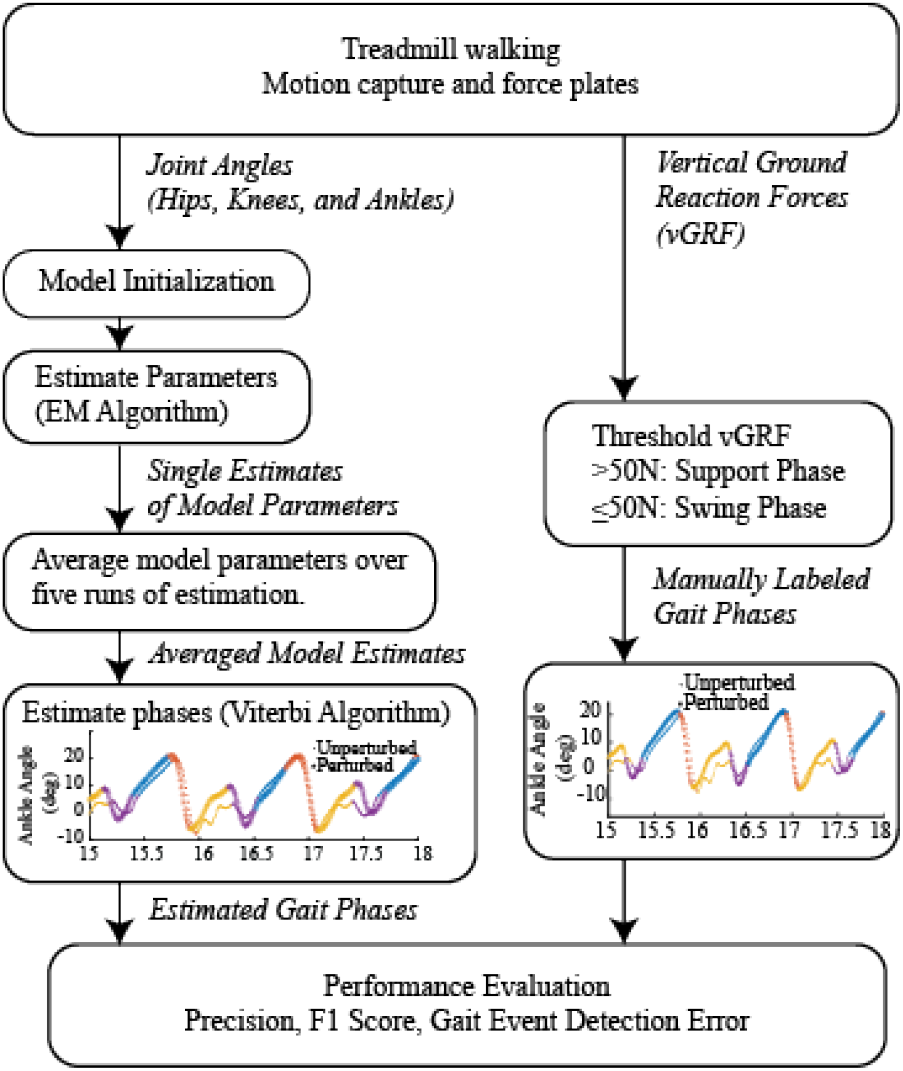
Flow diagram of the proposed methods. For each individual, joint angle kinematics are modeled using the proposed SLDS approach and the true gait phase labels are determined from ground reaction forces. Model parameters are averaged over five separate estimates and used to calculate the model-inferred gait phases via the Viterbi algorithm. Estimated phases are compared to the manually determined gait phases to evaluate the performance of the model. Inserts show the results of SLDS gait phase labeling and manual labeling on gait for the right ankle flexion in both unperturbed and perturbed walking

### D. Verification

Using force plate data, we constructed a ground-truth gait phase sequence to compare our models predictions against. In keeping with other gait event-detection studies [14,15], we thresholded ground reaction forces to define our ground-truth gait events. For each leg, heel-strikes were defined as when the force first exceeds 50N, while toe-offs were defined as when the force first drops below 50N. From the measured gait events, we generated a ground-truth gait phase sequence to label each point in our kinematic data.

Following training on baseline walking data, we used the SLDS models to infer the gait phases associated with each time point in baseline walking. We calculated the model fitness for each joint angle as one minus the normalized root mean squared deviation of the one-step ahead prediction residual:

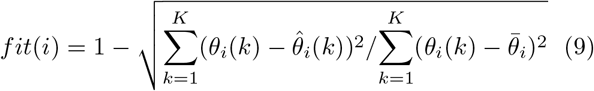

where *θ_i_* is the measured value, 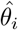 is the predicted value, and 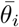 is the average value of the *i^th^* joint angle. From the hidden states and measured gait phases, we calculated the accuracy, precision, recall, and F1 score of the model for each phase. We treated each gait phase as a one-vs-rest classification problem, and calculated the true positive (TP), true negative (TN), false positive (FP), and false negative rate (FN). Then, the performance metrics for each gait phase were calculated as:

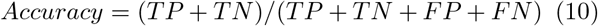

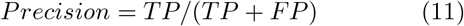

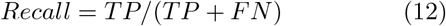

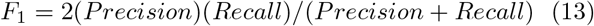

We further tested the ability of the SLDS to model joint angle kinematics and to identify gait phases during perturbed gait. Across individuals, we compared the model fitness for each joint angle in perturbed walking to the model fitness for baseline walking using a paired, one-tailed t-test. We also calculated the accuracy, precision, recall, and F1 score of the model for each gait phase on perturbed walking to baseline walking using a paired, one tailed t-test. We compared the gait event times of the model to ground truth in both normal and perturbed walking. We defined the timing error as the difference between the model transitions and nearest corresponding ground-truth event. For each individual in both baseline and perturbed walking, we calculated the average difference between the model transitions and the measured gait events for left heel-strike (LHS), right toe-off (RTO), right heel-strike (RHS), and left toe-off (LTO).

### E. Comparing SLDS with previous methods

Previous work using GHMMs to identify gait phases has not reported performance measures based on ground truth labeling of the gait phases. To compare our model against GHMMs, we trained a four-state GHMM for each individual in baseline walking, and calculated the GHMM-inferred gait phases using the Viterbi algorithm. We calculated the accuracy, precision, recall, and F1 score of the GHMM for each individual in both baseline and perturbed walking and compared the performance to that of our SLDS models.

## III. Results

### A. Four-state SLDS can represent joint angle trajectories

Across individuals, the SLDS models account for 88-99% of the deviation in each joint angle in baseline walking, with a median of 97% deviation accounted for across joint angles. The average SLDS fitness across individuals in normal walking was greater than 0.90 for all joint angles. In perturbed walking, the average fitness across individuals was greater than 0.90 for all joint angles except right ankle flexion (Table 2). The fitness of the SLDS was significantly less (p<0.05) for perturbed walking compared to normal walking in several joint angles, with most of significant differences occurring in the joint angles of the right leg.

**Table II.** SLDS Reconstruction Fitness

| Joint Angle | Reconstruction Fitness |  |  |
| --- | --- | --- | --- |
|  | Baseline Walking | Perturbed Walking | p-value |
| Left Hip Flexion | 0.99 ± 0.00 | 0.98 ± 0.01 | 0.13 |
| Left Hip Adduction | 0.95 ± 0.02 | 0.94 ± 0.03 | 0.15 |
| Left Knee Flexion | 0.98 ± 0.01 | 0.97 ± 0.03 | 0.10 |
| Left Ankle Flexion | 0.95 ± 0.02 | 0.92 ± 0.04 | 0.04 |
| Left Ankle Adduction | 0.93 ± 0.04 | 0.91 ± 0.05 | 0.05 |
| Right Hip Flexion | 0.99 ± 0.00 | 0.98 ± 0.01 | 0.01 |
| Right Hip Adduction | 0.95 ± 0.03 | 0.90 ± 0.07 | 0.08 |
| Right Knee Flexion | 0.99 ± 0.01 | 0.98 ± 0.01 | 0.03 |
| Right Ankle Flexion | 0.94 ± 0.02 | 0.86 ± 0.08 | 0.03 |
| Right Ankle Adduction | 0.96 ± 0.01 | 0.93 ± 0.02 | 0.06 |

### B. SLDS hidden states robustly correspond to gait phases

Across individuals, the hidden states of the SLDS correspond to the gait phases with 79-90% precision for baseline walking, averaged across the gait phases (Table 3). In a representative example from one individual (Figure 1), kinematically similar postures are grouped together by the model-inferred gait phase labels, indicating the model hidden states correspond to contiguous regions in state space. Following training, each of the linear dynamical models in the SLDS represents the evolution of the joint angle kinematics within each of the gait phases.

**Table III.** Average Precision for SLDS and GHMM by Participant

| Participant | SLDS |  | GHMM |  |
| --- | --- | --- | --- | --- |
|  | Baseline | Perturbed | Baseline | Perturbed |
| 1 | 79 ± 23% | 71 ± 25% | 86 ± 13% | 86 ± 20% |
| 2 | 84 ± 17% | 90 ± 2% | 66 ± 15% | 73 ± 16% |
| 3 | 86 ± 8% | 84 ± 13% | 90 ± 11% | 85 ± 17% |
| 4 | 89 ± 13% | 87 ± 14% | 87 ± 15% | 82 ± 20% |
| 5 | 85 ± 15% | 86 ± 18% | 84 ± 17% | 87 ± 16% |
| Grand Mean | 85% | 84% | 82% | 82% |

Even with the perturbations to gait kinematics from the muscle stimulation, the SLDS model for normal walking data still precisely recognized the gait phases with no significant difference in precision across participants (paired t-test, p>0.05). Averaging across all the individual-specific models, the SLDS identifies the gait phases with 85% precision, using only the dynamics learned from baseline walking. The average F1 score for each gait phase ranged from 79-89% on the perturbed data; moreover, the F1 score on perturbed walking was within 3% of the F1 score for baseline walking for each gait phase (Table 4).

**Table IV.** Average Performance for SLDS and GHMM by Gait Phase

|  | F1 Score by Gait Phase |  |  |  | Average F1 | Precision | Recall | Accuracy |
| --- | --- | --- | --- | --- | --- | --- | --- | --- |
|  | Left Swing | Left Double Support | Right Swing | Right Double Support |  |  |  |  |
| SLDS on Baseline | 87 ± 6% | 82 ± 5% | 88 ± 11% | 80 ± 9% | 84 ± 8% | 85 ± 15% | 87 ± 13% | 93 ± 4% |
| SLDS on Perturbed | 89 ± 5% | 80 ± 6% | 86 ± 16% | 79 ± 15% | 83 ± 11% | 84 ± 16% | 86 ± 13% | 92 ± 5% |
| GHMM on Baseline | 87 ± 8% | 80 ± 9% | 87 ± 9% | 79 ± 14% | 83 ± 10% | 82 ± 16% | 86 ± 12% | 92 ± 5% |
| GHMM on Perturbed | 85 ± 7% | 80 ± 7% | 90 ± 4% | 78 ± 11% | 83 ± 8% | 82 ± 17% | 87 ± 11% | 92 ± 3% |

### C. Transitions between SLDS states correspond to independently measured gait events

For four of the five individuals, transitions between SLDS hidden states matched the timing of gait events measured by force plates within a maximum average difference of 85ms for baseline walking. In the fifth individual (participant 1), the average difference for detecting right heel strikes was 167ms, while the average differences for the remaining gait events was <30m. While the difference in time varied across gait cycles, with the model predictions leading at some instances (negative difference) and lagging at others (positive difference), the transitions between LDS models matched the measured gait events on the order of 10s of milliseconds, with a median absolute difference of 25ms across individuals and gait events. An example from one participant illustrates the trend that toe-offs are identified early by the SLDS, and heel-strikes are identified late (Figure 3). Furthermore, similar time differences were measured in the perturbed walking case, where for four participants the maximum average difference was 97ms and the median absolute difference was 27ms. In the fifth individual (participant 1), the maximum was 143ms for right toe off.

**Fig. 3.**
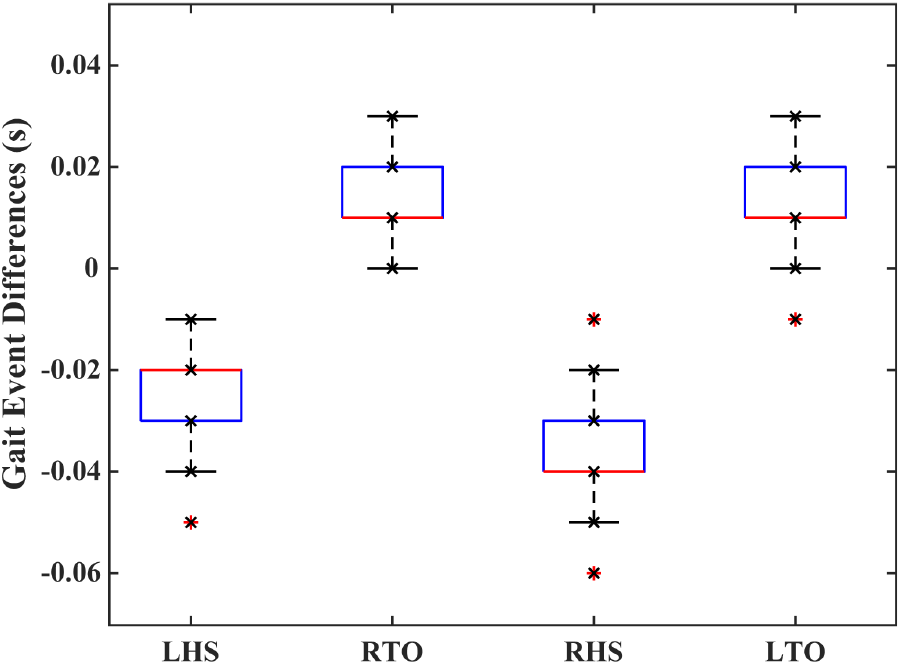
Difference between times of ground-truth gait events and model-inferred gait events for left heel-strikes (LHS), right toe-offs (RTO), right heel-strikes (RHS), and left toe-offs (LTO). A negative difference indicates the model-determined gait event occurs before the force-plate measured gait events. The figure shown is a sample from one individual in the normal walking condition.

### D. SLDS identifies gait phases with similar performance to GHMM

In three of the five participants, SLDS models identified gait phases with <5% difference in precision compared to GHMM models. In one participant, the SLDS outperformed the GHMM by >10% average precision in both baseline and perturbed walking, while in another participant, the GHMM outperformed the SLDS by 7% in average precision for both baseline and perturbed walking (Table 3). In individual gait phases, there was <5% difference in average F1 score between SLDS and GHMM for all phases in both baseline and perturbed walking data (Table 4).

## IV. Discussion

We demonstrated that SLDS models of gait can be used to infer gait phases and gait events from joint kinematics, even when gait dynamics are perturbed. We modeled discrete changes in lower limb dynamics using individual-specific SLDSs, with each LDS representing inter- and intra-limb dynamics in each of four gait phases. Switching times for the LDSs corresponded to actual gait events measured independently through ground-reaction forces and were not affected by perturbations that altered interjoint kinematics during gait. Because the SLDS explicitly represents gait dynamics and can be computed in real time, it may be a useful way to model gait for HRI aimed at improving gait.

Here, we demonstrated that the entire training procedure for the SLDS can be achieved with kinematic data only, given appropriate initialization. Initialization methods are seldom described in the literature, but are necessary for learning models of data with defined transitions, such as gait. With a random initialization, we found that model training can result in hidden states and dynamical models with no apparent meaning. Our initialization procedure relies on kinematic features of the data, and biases each LDS towards learning the dynamics of a single gait phase. We also constrained the hidden state transitions such that each state can only lead to one other hidden state, resulting in a cyclic pattern to mimic gait. The combination of the structured initialization and state transition constraints provided our models with enough domain knowledge to learn the gait phases and dynamics solely from the available joint kinematics during training.

Our work demonstrating the feasibility of SLDS on simple gait is a first step towards using SLDS to represent more complex gait behaviors. We showed that one LDS per hidden state was sufficient to model nominal gait at a fixed speed. However, we also demonstrated that one LDS per hidden state will not account for all the variation in novel gait types, as evidenced by the decrease in model fitness in the perturbed gait case. A bank of linear systems associated with each phase could describe a set of possible gait dynamics for each gait phase [8]. The individual LDS models would describe a gait style within a gait phase, such as nominal swing or swing with functional electrical stimulation. Switching between different systems in the bank over time could be useful for modeling and understanding complex gait variations, but would require large amounts of data to train.

To our knowledge, this is the first time a machine learning model of gait phase based on joint angle kinematics has been validated by an external measure of gait, instead of against pre-defined gait phase durations. We validated the SLDS-inferred gait phases by comparing them to gait phases determined from a force plate and demonstrated that SLDS precisely inferred gait phase from joint kinematics. Previous work with SLDS models of joint angle kinematics assumed the individual LDS models correspond to gait phases [6–8]. More recently, models of gait phase based on ground reaction forces have been proposed and validated against independent measures of gait phase [14]. These models use multiple regression HMMs (MRHMM) to infer seven phases of gait from vertical ground reaction force. Across all gait phases, our SLDS models achieved the same performance as the MRHMMs for identifying gait phases. Comparing our results to previous work [14] indicates that models of joint angle kinematics may be as appropriate as models of ground reaction forces for determining gait phase. SLDS models may also be useful for estimating gait events in real-time when force data are unavailable or impractical.

We also validated our model by comparing the SLDS hidden state transitions to measured gait events. Many of the transition times between SLDS states were within 100ms of the measured gait events. Kinematic methods for determined gait events typically estimate events from local minima and maxima in the joint angles [15,16], and have been validated against a force plate. These methods have detected heel-strikes and toe-offs with the same average time difference as our SLDS approach. Other approaches using instrumented shoes have applied fuzzy logic and supervisory rules [17] or variational inference [18] to detected gait events, and have achieved similar average errors. While our proposed SLDS approach produces similar results, a direct comparison is difficult because differences in walking speed and sampling rate across studies alters the relative importance of millisecond accuracy. Future work would benefit from reporting gait event detection errors as percentages of the gait cycle, facilitating comparison across study conditions.

Our work demonstrating that SLDS hidden states correspond to gait phases is a first step towards using SLDS to predict future kinematic trajectories. One advantage of the SLDS approach over GHMMs is that the SLDS incorporates a dynamic model of the kinematics, while the GHMM models only the statistics of the joint angles in each phase. While here we validated the SLDS-inferred gait phases, we did not test the ability of SLDS to generate new gait kinematics. Our SLDS models assume that the gait phase at the next time point depends only on the current gait phase, and not on the current kinematic state. This assumption results in simulated gaits that switch between phases randomly, instead of when the foot makes or breaks contact with the ground. Prior work has relied on knowledge about the SLDS switching times to generate recognizable gaits [6,8]; however, modeling the dependence of gait phase on gait kinematics could be more useful for modeling gaits with highly variable or undetermined gait phase durations.

We have demonstrated that SLDS models of gait incorporate an explicit model of gait phase as well as interjoint dynamics for single speed walking. Our SLDS, however, modeled only a single speed of treadmill walking in healthy young adults. In an HRI setting, gait speed will vary during overground walking, and the complexity of gait dynamics may vary with gait impairment. currently, the ability of SLDS to capture gait dynamics and model gait phases under these conditions is unknown. While our results may be suitable for use in treadmill walking scenarios, more work is necessary to allow the SLDS to account for gait variability.

## V. Conclusions

Switched linear dynamical systems can provide individual-specific representations of gait that incorporate both gait phases and gait dynamics. Our SLDS models identify gait phases from kinematics with the same precision as GHMMs, and have comparable performance to unsupervised models of ground reaction forces. Because the hidden states correspond to gait phases, each of the linear subsystems represent linearized lower limb dynamics within each phase. Our work here is the first step towards developing individual-specific models of gait dynamics for HRI targeting gait rehabilitation. Future work modeling the external effects on gait with SLDS could allow for HRI control strategies to be specifically designed for each gait phase.

## Acknowledgment

The authors thank Dr. Jessica Allen of West Virginia University and Dr. Trisha Kesar of Emory University for providing the data used in this study.

